# Flotillin-Associated rhodopsin (FArhodopsin), a widespread paralog of proteorhodopsin in aquatic bacteria with streamlined genomes

**DOI:** 10.1101/2023.01.04.522823

**Authors:** Jose M. Haro-Moreno, Mario López-Pérez, Alexey Alekseev, Elizaveta Podoliak, Kirill Kovalev, Valentin Gordeliy, Ramunas Stepanauskas, Francisco Rodriguez-Valera

## Abstract

Microbial rhodopsins are often found more than once in a single genome (paralogs) that often have different functions. We screened a large dataset of open ocean single-amplified genomes (SAGs) for co-occurrences of multiple rhodopsin genes. Many such cases were found among Pelagibacterales (SAR11), HIMB59 and the Gammaproteobacteria *Pseudothioglobus* SAGs. These genomes always had a *bona fide* proteorhodopsin and a separate cluster of genes containing a second rhodopsin associated with a predicted flotillin coding gene and have thus been named flotillin-associated rhodopsins (FArhodopsins). They are quite divergent from the other proteorhodopsin paralog and contain either DTT, DTL or DNI motives in their key functional amino acids. FArhodopsins are mainly associated with the lower layers of the epipelagic zone. All marine FArhodopsins had the retinal binding lysine, but we found their relatives in freshwater metagenomes that lack this key amino acid. Alfa-fold predictions of marine FArhodopsins indicate that their retinal pocket might be very reduced or absent, hinting that they are retinal-less (blind). Freshwater FArhodopsins were more diverse than marine FArhodopsins, but we could not determine if they are present as paralogs of other rhodopsins, due to the lack of SAGs or isolates. Although the function of FArhodopsins could not be established, their conserved genomic context indicated involvement in the formation of membrane microdomains. The conservation of FArhodopsins in diverse and globally abundant microorganisms suggests that they may be important in the adaptation to the twilight zone of aquatic environments.

**IMPORTANCE:** Rhodopsins have been shown to play a key role in the ecology of aquatic microbes. Here we describe a group of widespread rhodopsins in aquatic microbes associated with dim light conditions. Their characteristic genomic context found in both marine and freshwater environments indicates a novel potential involvement in membrane microstructure that could be important for the function of the co-existing proteorhodopsin proton pumps. The absence or reduction of the retinal binding pocket points to drastically different physiology. In addition to their ecological importance, novel rhodopsins have biotechnological potential in the nascent field of optogenetics.

## INTRODUCTION

Rhodopsins are probably the most universal biological light-energy transducers. They are found in all the domains of life (eukaryotes, bacteria, and archaea) and also in viruses (1). It is now well established that microbial rhodopsins (like microbes themselves) are barely represented by culture collections. Most of their diversity is found in the vast reservoir of uncultivated microbes present in the ocean and other aquatic environments exposed to light (2–6). Microbial rhodopsins were initially discovered in the halophilic archaeon *Halobacterium salinarum* (7), which contains four paralogous rhodopsin genes coding for different functions: H^+^ or Cl^-^ pumping, or light sensing with two different light ranges (8). In haloarchaea, the most common case is that more than one rhodopsin gene is found per genome reaching up to six described for *Haloarcula marismortui* (9). In other aquatic bacteria, the presence of multiple rhodopsin genes (paralogs) in the genome is much rarer, and only a few marine bacteria, such as *Dokdonia eikasta, Gillisia limnaea*, or *Nonlabens marinus* encode for, in addition to the regular proton-pumping rhodopsin, a sodium-pump and a chloride-pump (10–12). One potential reason for such scarcity is that many pelagic aquatic microbes have streamlined genomes that are depleted in paralog genes. Besides, they tend to be hard to grow and are largely known by metagenomic assemblies that are often incomplete. However, the presence of paralog copies of rhodopsins is interesting, particularly when the sequence is widely divergent since they may provide different functions to the microbe. Hence paralog rhodopsins are a potential source of novel functions of these versatile proteins that have also biotechnological potential for their use in optogenetics (13, 14).

Here we evaluate the genetic repertoire of rhodopsins in a large collection of single-amplified genomes (SAGs), which were generated from a global set of seawater samples from the tropical and subtropical, epipelagic ocean (15). The screening for rhodopsins resulted in approximately 3% of SAGs containing two type-I rhodopsin paralogs within their genomes. While one of them was predicted to be a standard proton pump, the other paralog belonged to a formerly described cluster of DTT-T rhodopsins discovered in contigs assembled from deep-water metagenomes in station ALOHA (16). The presence of this second rhodopsin was found restricted to three taxa known to have small and streamlined genomes-Pelagibacterales, HIMB59, and the Gammaproteobacteria *Pseudothioglobus*. This rhodopsin gene was always found next to a flotillin gene. Metagenomic screening in marine and freshwater samples revealed a broad distribution of these sequences in both types of environments, always associated with low light intensities. Our results on the protein structure, metagenomic recruitment, and genomic context can help to understand its function in these streamlined microbes.

## RESULTS AND DISCUSSION

### Paralog rhodopsin recovery and diversity

To identify rhodopsin paralogs, we first tried to locate them in the RefSeq database (April 2022) which includes cultivated marine microbes, but none was found, with the known exceptions mentioned above. This was not surprising considering the narrow range of diversity covered by cultivated marine bacteria, particularly of streamlined and abundant microbes. Fortunately, an alternative to monoclonal genomes of marine bacteria is single-amplified genomes (SAGs). Although many of them are incomplete they offer a significant guarantee that they start from a single cell and that the redundant genes found in them are real paralogs. Thus, we used a large collection of marine SAGs, composed of 12,715 partial genomes, which were produced using a random cell selection strategy from the epipelagic seawater samples from tropical and subtropical latitudes of the Atlantic and Pacific Oceans (15). The taxonomic composition of marine prokaryoplankton in this database is consistent with previous 16S rRNA amplicon and shotgun metagenomic studies of marine off-shore waters, indicating that this dataset is representative of the in situ microbial communities (15). We removed SAG assemblies with an estimated completeness lower than 50 %, which is the threshold for classifying them as medium quality (17). Finally, a total of 4,751 SAGs were screened for the presence of type-1 rhodopsin genes in their genomes (see methods). Among them, 63 % coded for type-I rhodopsins (**Figure 1A**). As previously described from metagenomic assemblies, most of these sequences were found in microbes from the order Pelagibacterales (#1,619), SAR86 (#386), Flavobacteriales (#258), HIMB59 clade (#191), and *Ca*. Actinomarinales (#139), as well as other less abundant taxa (**Table S1**). A small set of SAGs (∼3 %) coded for two type-I rhodopsins within their genomes. A close inspection revealed that they all belonged to three taxonomic groups: Pelagibacterales (SAR11), HIMB59 (formerly AEGEAN-169), and the gammaproteobacterial *Pseudothioglobus* **(Figure 1A)**, all typical marine streamlined microbes. To analyze the phylogenetic diversity of the recovered paralogs of type-I microbial rhodopsins, a maximum likelihood phylogenetic tree was constructed, including several reference sequences covering all types of rhodopsins. The phylogenetic tree showed that, although one of the rhodopsin sequences clustered according to their taxonomic classification and into the proton-pump proteorhodopsin family (**Figure 1B**), the sequences of the second rhodopsin paralog, which we named FArhodopsin, clustered on an independent branch (**Figure 1B**).

**Figure 1.**
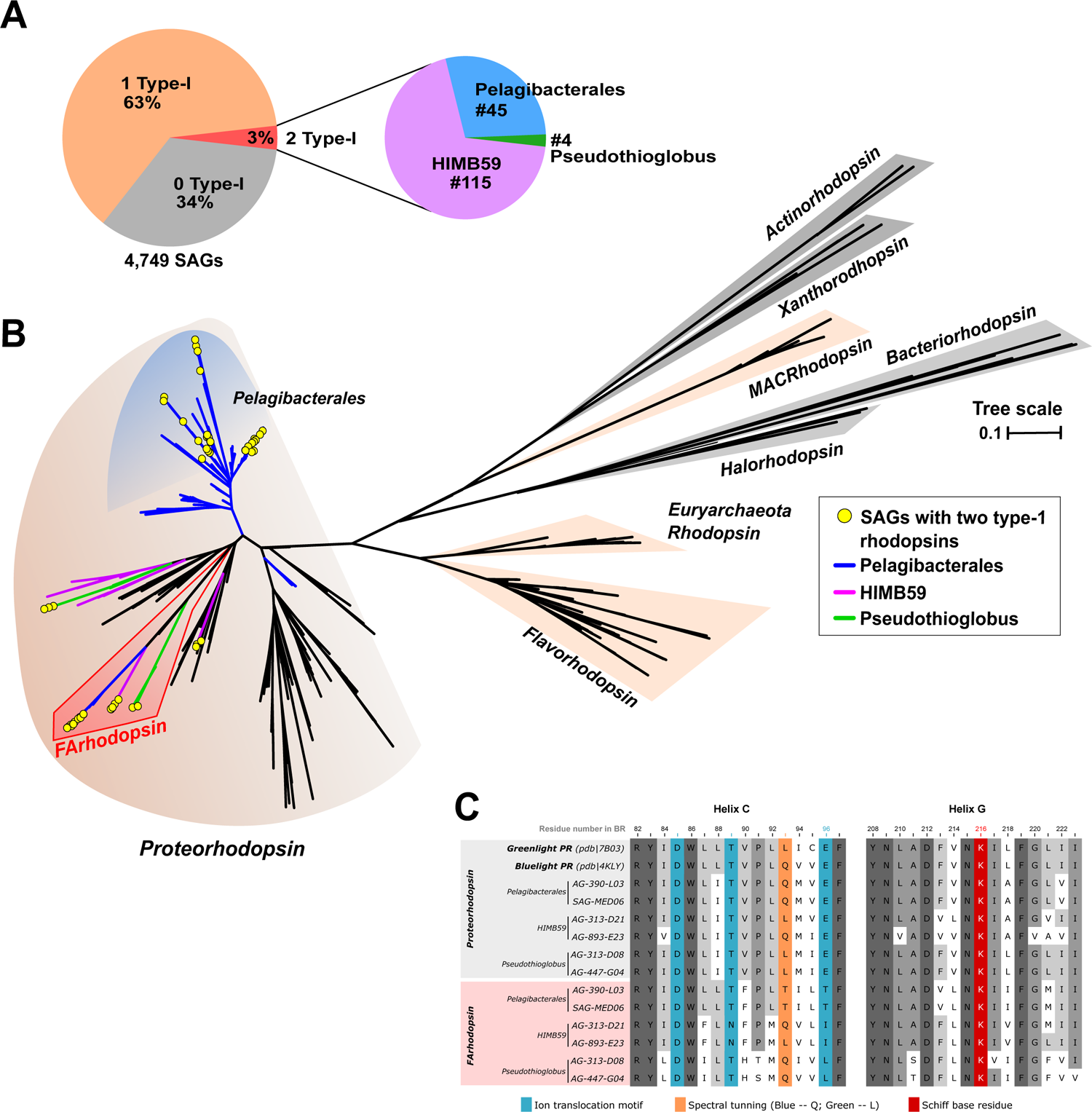
**A**. Pie charts indicating the percentage of single-amplified genomes (SAGs) in which two, one or no type-I rhodopsins were detected. Only those SAGs (from a large database collected from the tropical and subtropical euphotic ocean, Pachiadaki et al., 2019) that met the established quality criteria of ≥50% completeness and ≤5% contamination, i.e., medium to high quality draft genomes were included in the analysis. For those SAGs containing 2 rhodopsin genes, a second pie chart indicates their taxonomic affiliation **B**. Maximum likelihood phylogenetic tree of rhodopsin genes detected in the marine SAGs, together with several rhodopsin references covering all types of microbial rhodopsins. Branches have been colored for SAGs harboring two rhodopsins according to their taxonomy (blue, Pelagibacterales; purple, HIMB59 and green, *Pseudothioglobus*). Sequences belonging to the new clade of rhodopsins (Flotillin-associated rhodopsin; FArhodopsin) are highlighted in red. Branches within color background contain type-I rhodopsins from the SAG dataset. The yellow circles correspond to the SAGs containing the two type-I rhodopsin paralogs. **C)** Protein sequence alignment of Helix C and G residues for the two type I rhodopsins. Two reference proteorhodopsin proton pumps at the top. Residues involved in rhodopsin function (e.g. ion translocation) and wavelength absorption are highlighted in blue and orange, respectively. The lysine (K) highlighted in red (helix G) is responsible for the covalent binding of retinal. Dark and light grey regions represent conserved residues within the alignment.

An analysis of the protein alignment of the two paralog rhodopsins (**Figure 1C)** indicated that all seemed to be potentially functional. Both paralogs code for the lysine (K) in helix G responsible for the binding of the retinal to the protein via a Schiff base reaction although structural and experimental predictions make it unlikely that one of them bind retinal (see below). In addition to many major differences, including a deletion of 23 aa at the N terminus of the second rhodopsin (FArhodopsin), which might include a leader peptide **(Figure S1)**, we observed differences in the ion-pumping motif. While one of the paralogs showed the DTE motif (positions 85, 89, and 96 according to the amino acid sequence of bacteriorhodopsin from *Halobacterium salinarum*) identical to the green and blue light proton-pumping proteorhodopsins (**Figure 1C**), in the FArhodopsin we detected three variations. In the case of Pelagibacterales and the *Pseudothioglobus*, the FArhodopsin differed in the last amino acid of the triad, being threonine (DTT) as has been described previously for metagenomic assembled rhodopsins from ALOHA (16) and leucine (DTL), respectively, while in HIMB59, the DTE motif was replaced by aspartate-asparagine-isoleucine (DNI). In addition, the amino acid residue suggested being critical to the spectral tuning of rhodopsins (position 93 in bacteriorhodopsin) also changed. Leucine, methionine, and isoleucine residues are detected commonly in proteorhodopsins absorbing in the green light spectrum, whereas a rhodopsin containing glutamine absorbs in the blue light range. The FArhodopsin sequences of HIMB59 and *Pseudothioglobus* had either the glutamine or leucine, while the Pelagibacterales FArhodopsin coded for threonine at that residue.

### The genomic context of FArhodopsins

The genomic comparison of Pelagibacterales SAGs containing the second rhodopsin against the genomes of isolates HIMB083 (genomospecies Ia.3/V), HTCC7011 (genomospecies Ia.3/I), and HTCC1062 (genomospecies Ia.1/I) (18) indicated that FArhodopsins were always located on a gene cluster of approx. 26 Kb that was absent from the isolate genomes (**Figure 2A**). Although SAGs containing this cluster were phylogenetically distant from each other, as they belonged to different subclades (listed in **Figure 2B**), most of the cluster genes were conserved and syntenic (**Figure 2B**).

**Figure 2.**
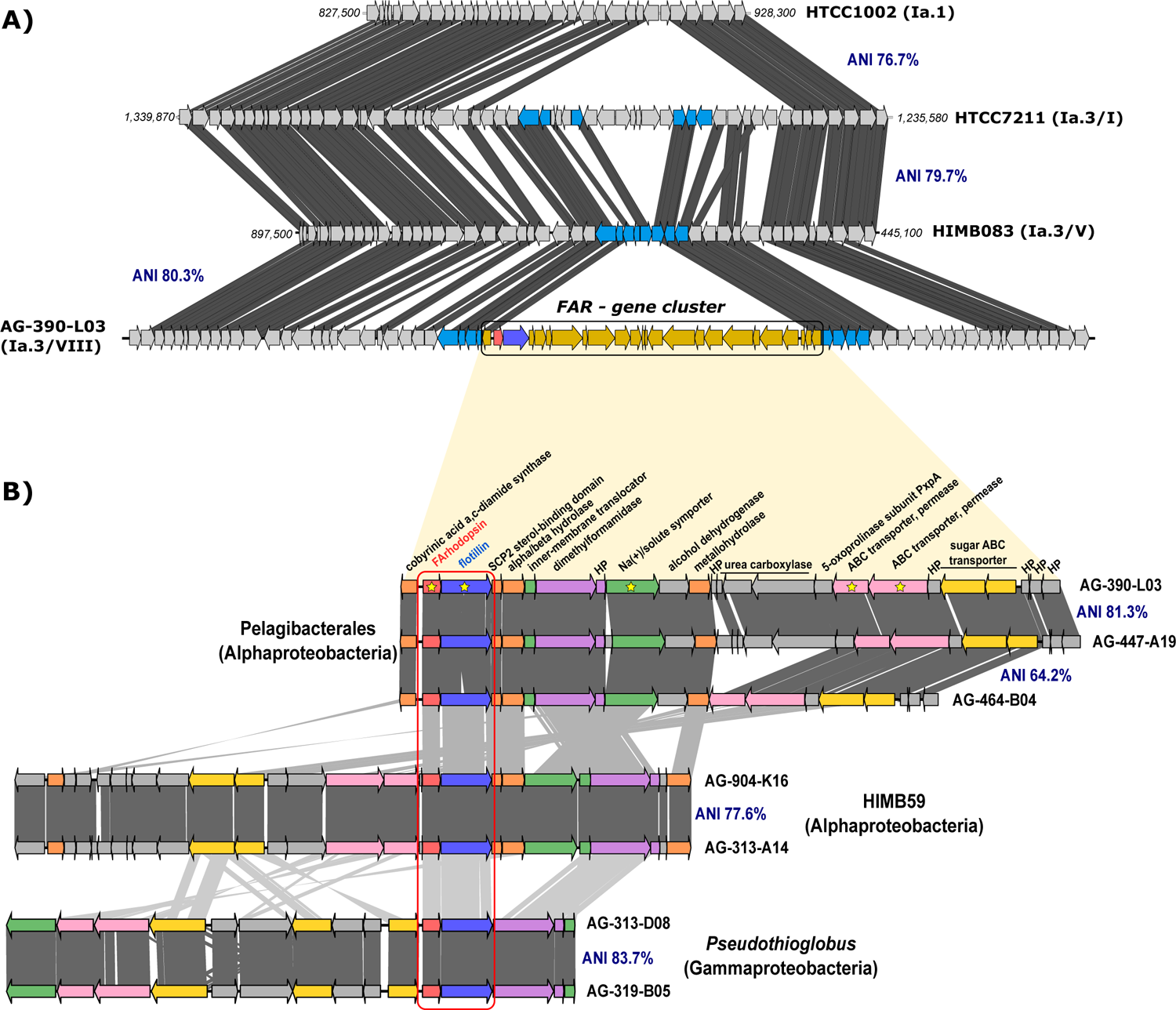
**A)** Genomic location of the FArhodopsin and its neighboring genes of Pelagibacterales SAG AG-390-L03 in relation to the reference genomes HTCC1002, HTCC7211, and HIMB083. Genes colored in yellow in the upper alignment represent the (flotillin-associated (FAR) gene cluster. The FArhodopsin and the flotillin are colored in red and blue, respectively. **B)** Genomic alignment (amino acids) of the flotillin-associated rhodopsin gene cluster found in Pelagibacterales, HIMB59 and *Pseudothioglobus* genomes. The ANI value for whole genome is indicated. The region containing the FArhodopsin and flotillin genes highlighted in a red box. Genes that are homologous are colored with the same color. Annotation is provided for the AG-390-L03 genome.

Despite some genomic rearrangements, the rhodopsin in the cluster always appeared co-localized on the same strand next (16 nucleotides intergenic spacer) to a large gene (1,977 bp) predicted to code for a flotillin. Flotillins are members of a widely conserved superfamily of proteins termed Stomatin, Prohibitin, Flotillin, and HfK/C (SPFH) domain proteins (19). Eukaryotic flotillins have been extensively studied (20) and only later they were found in prokaryotes (21, 22). They are classified as scaffold proteins associated with lipid rafts, lipid microdomains in membranes, which enhance the assembly of raft-associated proteins (23, 24). The gene clusters containing the rhodopsin in HIMB59 and *Pseudothioglobus* were delimited by comparing the genomes against closely related (>90% average nucleotide identity [ANI]) SAGs (the pure culture HIMB59 available was too distant). Regardless of the gain or loss of some genes, synteny was maintained among HIMB59 relatives, despite a wide phylogenetic distance. Even *Pseudothioglobus* (a gammaproteobacterium) genomes conserved the flotillin and the three last genes of the cluster located on the same strand and potentially co-transcribed (**Figure 2B and Table S2**). We evaluated the presence of flotillin proteins as well as any other “SPFH” domain-containing proteins in the initial set of marine SAGs (see methods). Among them, nearly 85% were positive for at least one “SPFH” domain-containing hit. A manual inspection resulted in a broad classification of these proteins as HflC and HflK. These proteins form the complex HflKC, involved in the regulation of the protease FtsH not related to the eukaryotic flotillins (25). Otherwise, the flotillin gene was detected mainly in genomes coding for the FArhodopsin (only twelve bona-fide flotillin proteins were detected not belonging to the rhodopsin gene cluster) (**Figure S2**). Phylogenetic classification of detected flotillins against well-characterized eukaryotic and prokaryotic sequences resulted in flotillin homologs from Pelagibacterales, HIMB59, and *Pseudothioglobus* forming a separate cluster (**Figure S2**).

The analysis of the genes found in the FArhodopsin cluster (FAR cluster) of the Pelagibacterales SAGs (**Figure 2B**) shows very conserved synteny despite the divergence (ANI) of the overall genomes. The gene cluster is always bordered by long spacers (236 and 219 nucleotides in the AG-390-L03) that are also well conserved. After the spacer, there is a set of ten genes on the same strand and likely also co-transcribed (see below). The next gene is annotated as a sterol-binding-domain of a sterol-carrier protein, likely involved in lipid transport. There is also an inner membrane translocator and a membrane sodium symporter. Thus, a correlation between membrane structure and function is strongly suggested. Only Pelagibacterales and HIMB59 encoded for a cluster of three genes annotated as biotin-dependent carboxyltransferases and carboxylases, that might be involved in the degradation of urea. Other metabolic genes detected included an N,N-dimethylformamidase, involved in the glyoxylate and dicarboxylate metabolism (26), the PxpA-like subunit, involved in the conversion of 5-oxoproline to L-glutamate (27), as well as some uncharacterized alpha/beta hydrolases, iron-dependent alcohol dehydrogenases, and aldolases. In addition, the *Pseudothioglobus* FAR cluster encoded for the transcriptional regulator LuxR and a hybrid sensor histidine kinase/response regulator (genes annotated in the FAR gene cluster are listed in **Table S2**).

### FArhodopsins in aquatic metagenomes

The presence of homologs to FArhodopsin in the ALOHA station depth profile has been extensively studied (16). The authors concluded that the DTT-T rhodopsin was present mainly in samples from 200 to 1000 m depth. Thus, to recover more genetic diversity of FArhodopsins we analyzed the assemblies of marine metagenomes, including western Mediterranean Sea depth profiles (28, 29), BATs (30), and global ocean surveys such as Tara Oceans (31) and GEOTRACES (30). A western Mediterranean sample has also been analyzed recently with long-read HiFi sequencing (32) that does not need assembly to recover complete gene clusters. Considering that microbes of the order Pelagibacterales, which are the main contributors of FArhodopsins, can be also found in freshwater ecosystems (e.g. *Fonsibacter*), we included in the analysis samples taken from lakes in North America (33), Europe, and Asia (34–36), including Lake Baikal (37, 38), where representatives of the *Pelagibacter* Ia.5 (18, 37) clade were recently found.

**Table S3** summarizes the number of rhodopsins type-I, type-III, FArhodopsins, and flotillins recovered in each metagenomic sample. In agreement with what we found in the dataset of marine SAGs, microbial type-I rhodopsins represented the majority of rhodopsin sequences in marine and freshwater assembled contigs, followed by heliorhodopsins. We could recover hundreds of FArhodopsin sequences, including from freshwater datasets (**Table S3**). This analysis showed the uneven recovery of FArhodopsin and flotillin sequences, which could call into question the strong association described previously between FArhodopsin and flotillin. However, a comprehensive study of the assembled contigs containing the rhodopsin revealed that most of them had FArhodopsins at the contig ends, indicating a breakpoint in the assembly, a common occurrence in regions of high microdiversity. Only in the metagenomic sample from the Mediterranean Sea sequenced with PacBio CCS (32), FArhodopsin and flotillin numbers were almost at par (407 and 455 sequences, respectively). All FArhodopsins well covered by the long reads contained the tandem confirming a consistent association. This result illustrates one of the major advantages of long-read metagenomics, allowing to recover complete genes and even operons from the bulk of CCS reads, without relying on metagenomic assembly.

The phylogeny of all of the recovered FArhodopsin proteins (#912) resulted in two major phylogenetic branches, separating the marine from the majority of the freshwater sequences (**Figure 3A**). All marine metagenomic FArhodopsin-containing assemblies were classified within Pelagibacterales, HIMB59, and *Pseudothioglobus* clades, regardless of depth, or geographical origin (e.g. latitude), indicating that apparently, no other microbes encode FArhodopsin globally. Interestingly, one cluster composed of sequences from deep samples of Lake Baikal (390 m, Russia) and Lake Constance (200 m, Switzerland) grouped within the Pelagibacterales FArhodopsin. A close inspection revealed a well-conserved synteny between these contigs and the marine SAG AG-390-L03 (**Figure 3B**). Based on gene annotation of the surrounding regions, these sequences belonged to the freshwater Ia.5 *Pelagibacter* clade, which was reported for the first time in freshwaters in Lake Baikal (37). Other freshwater metagenomes, on the other hand, showed a less restricted taxonomic distribution of FArhodopsins, with several distinct subclades indicating a higher sequence diversity (**Figure 3A**). Interestingly, protein alignment of these novel freshwater metagenomic sequences revealed that most did not encode for the lysine (K) in helix G responsible for the covalent binding of retinal to the protein, but they showed the amino acid variants glutamine, threonine, leucine, or arginine at this site (**Figure 3A and Table S4**). **Figure 3B** also shows that despite the divergence, sequences coming from freshwater preserve some genes of the FAR cluster found in the marine FARs. Besides, although the contigs harboring this protein were short hampering their taxonomic classification (as is typical of complex communities), those that were long enough always encoded for a flotillin next to the rhodopsin gene, highlighting the key role the flotillin gene plays in the cluster. Lysine-less rhodopsins (Rh-noK) have been already described for a group of xanthorhodopsins related sequences in marine betaproteobacteria (39) and were shown to be retinal binding and functional proton pumps when the lysine residue was reconstituted by genetic manipulation. Interestingly, Rh-noK was found located in tandem with a functional proteorhodopsin gene, and the authors speculated that both gene products could form a multimer in which Rh-noK could regulate the proton pumping activity of the neighbor (39).

**Figure 3.**
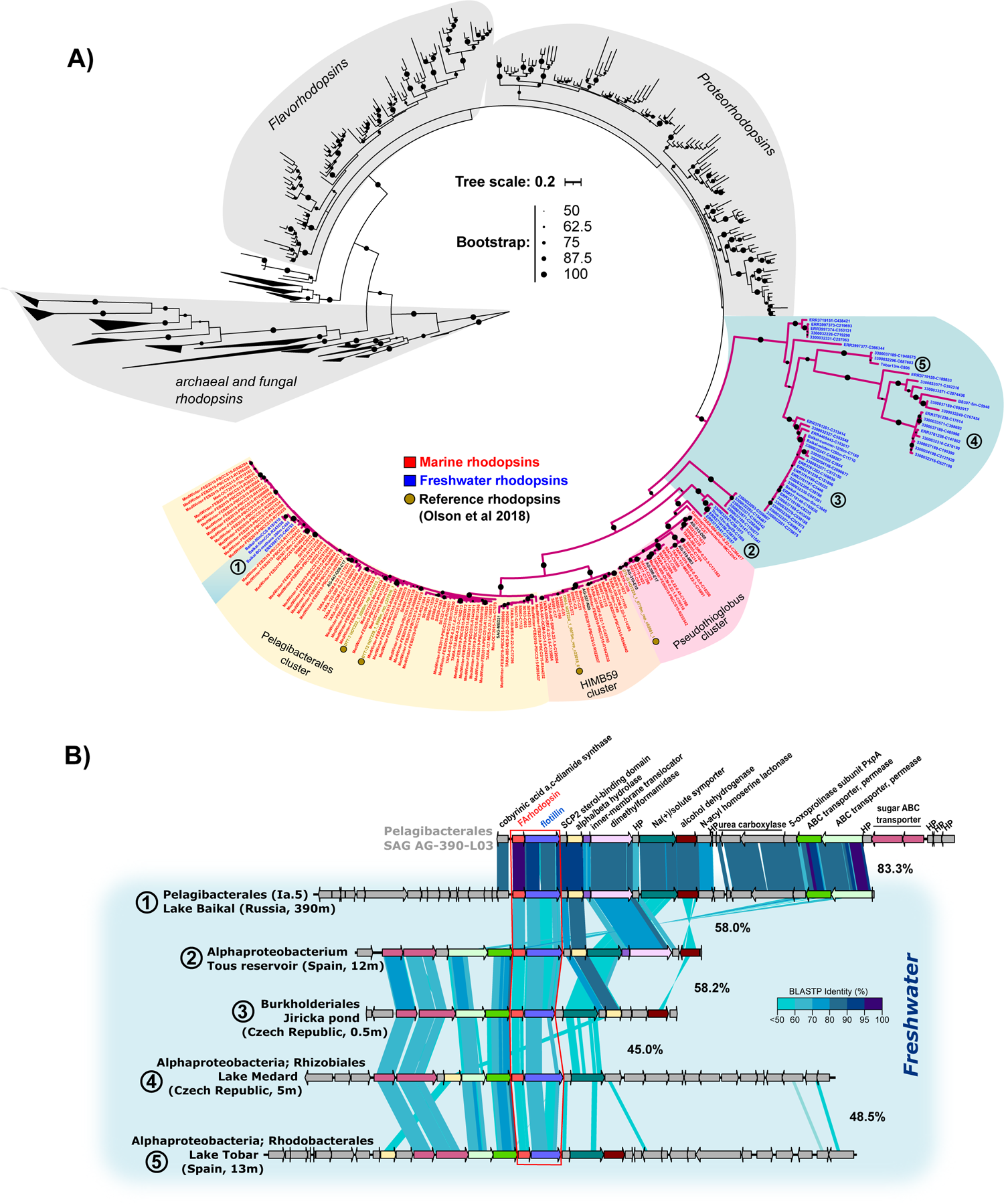
**A**. Maximum likelihood phylogenetic tree of FArhodopsin sequences detected from marine and freshwater datasets colored in red and blue, respectively. The full description of samples can be found in Table S4 and within Methods section. Sequences with 100% amino acid identity with those in the tree are not shown. Sequences in the blue background branch to the right (all from freshwater metagenomes) do not have lysine in the retinal binding position in helix G (Table S4). Numbers within freshwater branches represents clades on which a contig was used as reference for the genomic comparison shown in B. **B**. Genomic comparison of freshwater FArhodopsin subclades, using the Pelagibacterales SAG AG-390-L03 (subclade Ia.3) as a marine reference. Numbers between contigs represents the average amino acid identity (AAI, %) of shared proteins between them.

### Recruitment of metagenomic and metatranscriptomic fragments on the FAR gene cluster

To assess the abundance and distribution of this novel group of rhodopsins, we performed metagenomic fragment recruitment analysis by recruiting all the FAR gene clusters in the SAGs of Pelagibacterales (#24), HIMB59 (#69) and *Pseudothioglobus* (#10) taxa to 140 metagenomes from a depth profile in the Mediterranean Sea (29) and the *Tara* Oceans datasets (31). As expected, the three taxa showed differential depth recruitment in the Mediterranean Sea (**Figure 4A**) and the same was found for the *Tara* Oceans datasets containing deeper samples (**Figure S3**). Pelagibacterales are the most abundant microbes in surface layers of the water column, although it is known that this diverse order has an eurybathic distribution, with some clades being detected in the deep chlorophyll maximum (DCM), lower photic (LP) metagenomes, and in meso- and bathypelagic samples (29). However, the Pelagibacterales FAR cluster recruited more in deeper samples such as the DCM, lower photic and bathypelagic samples, represented by 45, 60, 75, 90, 1000, and 2000m depths in the available Mediterranean metagenomes (30) (**Figure 4A**). A similar preference for deeper waters was found globally with the *Tara* Ocean samples (**Figure S3**). Thus, confirming the trend discovered for the DTT-T rhodopsins found by Olson et al at ALOHA (16). In the Mediterranean metagenomes, the maximum recruitment of the cluster was found at the LP (75-90 m), much shallower than the 500 m detected at the more oligotrophic, permanently stratified waters at ALOHA (16). A similar trend was found for the Gammaproteobacteria *Pseudothioglobus*. Contrastingly, HIMB59 FAR (with the DNI motif in the FArhodopsin) recruited the most at the upper photic (15-30 m deep), following closely the recruitment of the full SAGs (data not shown) that were detected almost exclusively in the upper photic. In addition, we could not find a clear association of freshwater FArhodopsins with deeper waters, in Lake Baikal they appeared near the surface and down to 390 m deep but not in a bathypelagic sample (data not shown). In freshwaters, the photic zone is harder to delimit given the diversity of trophic status that characterize lakes and their overall shallower depths.

**Figure 4.**
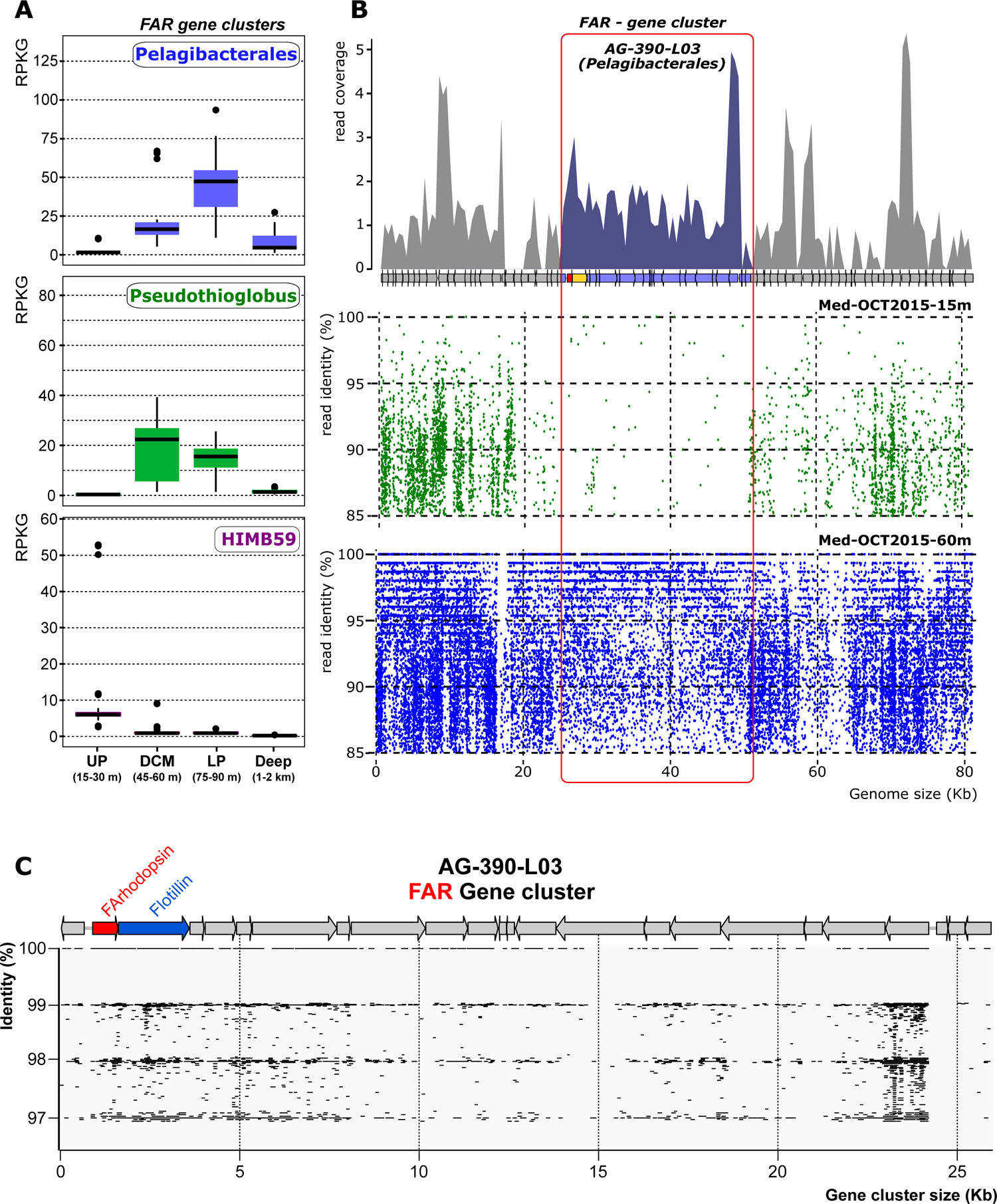
**A**. Relative abundance (measured in reads per kilobase of genome per gigabase of metagenome -- RPKG) of the flotillin-associated rhodopsin gene clusters in a depth profile metagenomic study from Western Mediterranean Sea collected in a single offshore location during a period of marked thermal stratification (Haro-Moreno *et al*. 2018). Metagenomes are grouped by depth (i) upper photic (UP, 15 and 30 m), (ii) deep chlorophyll maximum (DCM, 45 and 60 m), (iii) lower photic (LP, 75 and 90 m), and deep (Deep, 1,000 and 2,000 m) layers. **B**. Metagenome and Metatranscriptome analysis of flotillin-associated rhodopsin gene cluster. The upper panel show read coverage of the metatranscriptomic Illumina reads from the 60 m sample of the profile (98% identity, 250 bp window). In the lower panel, a metagenomic recruitment plot using raw reads from the samples 15m and 60m (>70% identity, >50 bp long) is represented. FAR gene cluster is highlighted in red. **C**. Read coverage showing individual reads in the 60 m metatranscriptome of the FAR gene cluster showing that the FArhodopsin and flotillin genes are co-transcribed (there are reads covering both genes simultaneously).

Metatranscriptomes were available from the same seawater samples (Western Mediterranean Sea, 15 and 60 m) (40). Their recruitment on SAGs confirmed the transcription of the FAR cluster of Pelagibacterales AG-390-L03 at 60 m (**Figure 4B and 4C**) but, as expected, no matching transcripts were found at 15m. The tandem rhodopsin-flotillin was the second most recruited region in the cluster (**Figure 4B**), indicating that these genes are abundantly expressed. The average recruitment value (in reads per kilobase gene per million transcripts – RPKM) of the FAR (0.32 RPKM) was substantially above the average number of transcripts aligning to the genome (0.2 RPKM, with the rRNA ribosomal operon excluded). The transcriptomic read coverage indicates clearly that both FArhodopsin and flotillin (at least) are cotranscribed in a single polycistronic mRNA despite the 16 nucleotides separating both genes (**Figure 4C**). Several mRNA reads containing the spacer could be detected (data not shown).

### Predicted structure of the retinal binding pocket

The most notable characteristic of microbial rhodopsins is their capability to respond to light through the use of the retinal cofactor. However, when (16) expressed in *Escherichia coli* a rhodopsin gene containing the DTT motif, they detect no binding of retinal, judging from the pellet pigmentation of centrifuged cells nor change in pH upon illumination. The lack of function or pigmentation in *E. coli* can be associated with a lack of proper membrane insertion into a lipid raft due to the lack of the flotillin-associated protein. Interestingly, Olson *et. al*. (16) modified a functional DTE-Q sequence to DTT-T, which resulted in the loss of function.

We used AlphaFold (41) to predict the structure of marine FArhodopsin to examine its retinal binding pocket. The predicted structure showed high similarity to that of the closest homolog with an experimentally determined high-resolution structure (PDB: 4jq6, blue proteorhodopsin from Med12, a predicted protein from a metagenomic sample from the Mediterranean Sea at 12 m deep (42)), which allowed us to adapt the retinal position to the predicted structure. However, when analyzing amino acids that are highly conserved among Pelagibacterales FArhodopsins (present in more than 99% of cases), we observed that four of these residues would intersect the retinal in its all-trans conformation (**Figure 5A**). A similar observation was made in other groups of marine FArhodopsins (**Figure 5B and 5C**). In addition, as mentioned above, most of the freshwater FArhodopsins lack the lysine that is used to bind retinal. These findings suggest that all FArhodopsins may be retinal-less like those previously described in the Halobacteriaceae (43) or betaproteobacteria Rh-noK (39). Their seven transmembrane helices may have evolved to perceive other stimuli, similar to G protein-coupled receptors (G-PCRs), which have adapted for responding to both light and chemical activation or they could be involved in either regulation or membrane microdomain organization as shown for some G-PCRs in the animal retina (44). However, it is important to note that there is no direct evidence that the AlphaFold prediction is precisely correct. Recently, an unwound helix in a structure of a channelrhodopsin called ChRmine was not predicted by AlphaFold (45). It is possible that another significant distortion from the classic rhodopsin fold could still allow for binding of the retinal, or that FArhodopsins may bind other carotenoid derivatives to absorb light.

**Figure 5.**
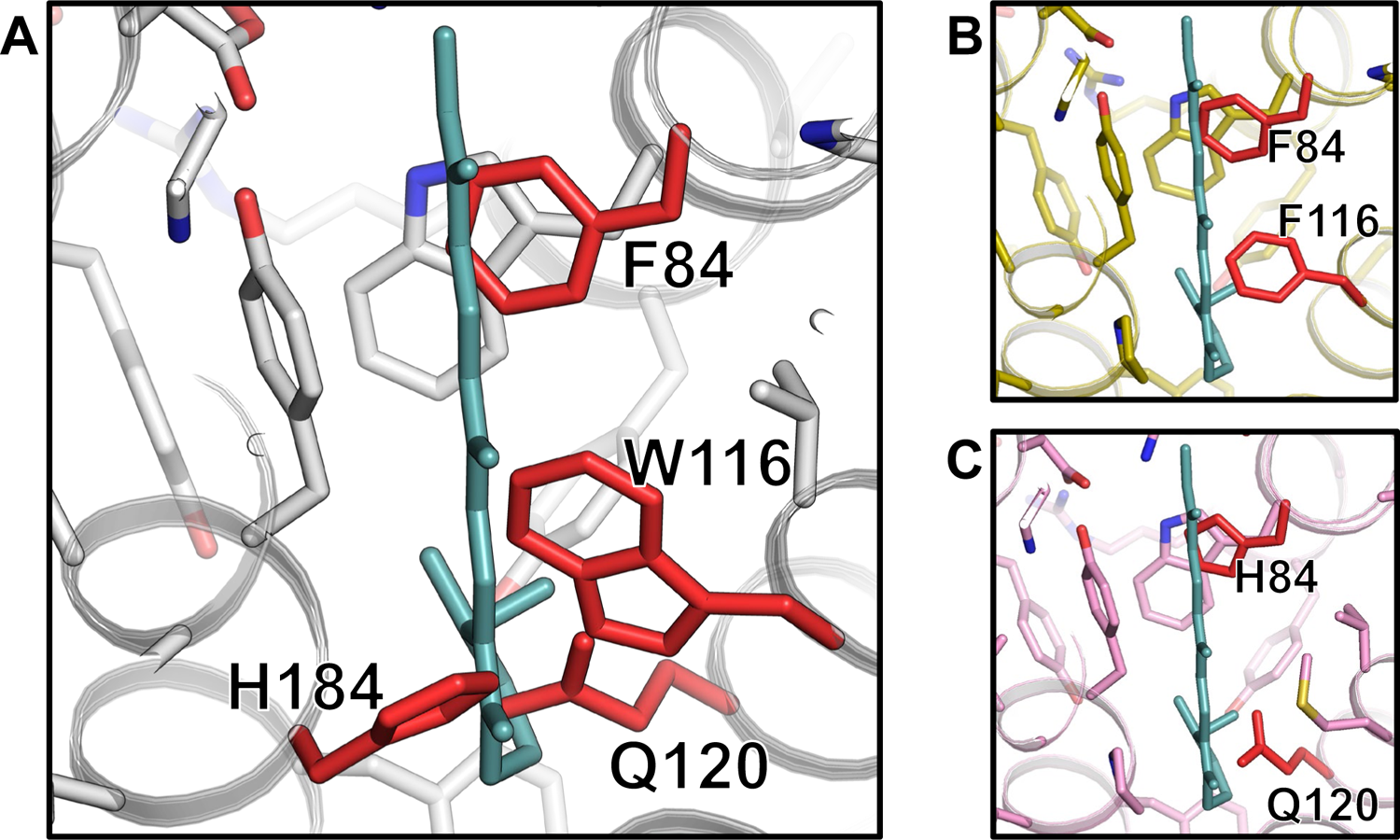
Alpha fold prediction of retinal binding pockets in marine FArhodopsins from different subclades: Pelagibacterales **(A)**, HIMB59 **(B)**, and *Pseudothioglobus* **(C)**. Retinal is depicted in aqua green, and only residues that are present in more than 99% of the subclade family members are displayed. Residues that clash with the retinal are indicated and highlighted in red.

### SYNTHESIS

The discovery, two decades ago (2), of rhodopsin sequences in marine prokaryoplankton communities resulted in a significant leap forward in the understanding of the biology of these microorganisms and their interaction with the ecosystem. In the photic zone, the proton-pumping activity of rhodopsins may provide “cheap” solar energy (46). However, rhodopsins can perform a wide variety of other functions (47, 48). Furthermore, the presence of paralogs with very divergent activities and even sequences has been known since the first microbial rhodopsins were described in halophilic archaea, and they are intensively looked after since they reveal novel physio-ecological roles and can be of potential application for optogenetics. In this work, we have analyzed the distribution and the presence of rhodopsin paralogs in a large dataset of marine SAGs (15) that provide strong evidence that the detected genes come from the same genome (15). We tried with the meager representation of cultured microbes but, unsurprisingly, could not find any examples of genomes with two rhodopsins. MAGs, on the other hand, are a composite of genomic fragments, and the presence of contigs belonging to different organisms (chimerism) may interfere with the analysis.

Unsurprisingly, only a small subset of SAGs coded for a rhodopsin paralog, most of which belonged to well-known streamlined lineages, such as Pelagibacterales (49–52), *Ca*. Actinomarinales (53, 54), and SAR86 (55, 56). These microbes are characterized by having a small genome size, a higher coding density, and a reduced set of paralogs, to cope with limited nutrient and energy availability in the oligotrophic open ocean (57). Nevertheless, we detected two copies of the rhodopsin gene in the genomes of three widespread streamlined taxa, Pelagibacterales, HIMB59 and the Gammaproteobacteria *Pseudothioglobus*. One paralog was a typical proteorhodopsin proton pump, while the other affiliated with a novel clade described among assembled contigs in ALOHA metagenomes at depths of 200 to 1000 m (16). The access to the genomic context, which was not possible previously, showed a consistent association with a conspicuous flotillin gene forming a single mRNA transcript. Flotillins are members of a superfamily of proteins termed “SPFH” (Stomatin, Prohibitin, Flotillin, and HfK/C) (19), detected for the first time in the cytoplasmic membrane of eukaryotes (20). These proteins are involved in the formation of microdomains (also known as lipid rafts) in the plasma membrane of all mammalian cells (20). Its biological role is not well understood, but it is believed that they act as scaffold proteins, enhancing the recruitment of raft-associated proteins to these microdomains to be active, as well as to facilitate their interaction and oligomerization with other proteins (23). Flotillin sequences have been also detected in bacteria. Specifically in *Bacillus subtilis*, which encodes for two subunits, YuaG and YqfA (renamed as FloT and FloA) (23). Mutants lacking both genes showed reduced sporulation efficiency and changes in the biofilm formation (21, 22, 58). Remarkably, our results indicated that the flotillin encoded in the FAR cluster was the only copy present in the genome, and therefore, its presence might be essential for the formation of specific microdomains in these organisms. In *B. subtilis* (and other bacteria) flotillin appears in lipid rafts that concentrate the phospholipid cardiolipin and cyclic (hopanoids) or non-cyclic isoprenoids, such as those derived from carotenoids (23). The genomes of Pelagibacterales, HIMB59, and *Pseudothioglobus* encode for a cardiolipin synthase. For that reason, the presence of lipid rafts, despite not being demonstrated “*in vivo*”, seems plausible. We would like to hypothesize that the flotillin detected in the genomic island might be critical for the function of the FAR cluster genes. Many FAR genes annotated were classified as membrane transporters or signal transduction/regulators, proteins typically enriched in flotillin-associated microdomains (21, 24).

In the Mediterranean, FArhodopsin genes were abundant in metagenomes coming from the lower photic zone, with a maximum at the deep-chlorophyll-maximum (45-60 m deep). In contrast, FArhodopsin genes peaked in deeper waters (500 m) at ALOHA station in the Pacific Ocean. This fact, together with the lack of *in vitro* activity and even retinal binding of some ALOHA representatives cloned in *E. coli* suggests that these rhodopsins might be retinal-less or even blind rhodopsins (not using light). Furthermore, most freshwater FArhodopsins do not have the lysine residue needed for retinal binding. However, a more in-depth screening of several metagenomic datasets indicates that they tend to associate with the photic (or twilight) zone, but not with bathypelagic waters. In addition, we have always detected a standard retinal-binding proton pumping proteorhodopsin present elsewhere in the genome, which indicates a clear light-interactive lifestyle (at least periodically). A possible explanation would be that flotillins enhance the formation of a heterocomplex of DTE and FArhodopsins in lipid rafts. This kind of rhodopsin heterocomplexes has been described in fungi (59, 60), on which a classic blue or green-absorbing rhodopsin cyclase (a kind of rhodopsin linked to a type III guanylyl cyclase) interacts with another rhodopsin cyclase, but with an absorption in the near-infrared spectrum. Thus, there is a chance that the FArhodopsin interacts with the proton-pump proteorhodopsin, present in the same genome, in a heterocomplex, modifying the spectral tunning to accommodate ion transport to deeper waters.

The presence of FArhodopsins in freshwater metagenomes came as a surprise. They are much more diverse at the sequence level than marine FArhodopsins and are present in a much wider taxonomic array of prokaryotes. The association with flotillin was also detected in all cases in which contigs were long enough suggesting similar functional properties. In one case, we found a FAR cluster syntenic with a marine *Pelagibacter* genome in a distant relative from Lake Baikal. The high similarity of the FAR cluster between marine and freshwater Pelagibacterales might indicate marine-to-freshwater transitions of the FAR gene cluster (37).

There is always the possibility that FArhodopsins act as sensors of other stimuli (as hypothesized for retinal-less rhodopsins of haloarchaea (43) or Rh-noK (39), but the lack of genes coding for transducers in the FAR cluster and the association with the photic zone in many metagenomes indicates otherwise. We hypothesize that FArhodopsins may act as complements to the *bona fide* proton pump, by increasing its performance under the dimmer light conditions prevalent in the lower photic zone. The streamlined nature of genomes containing FArhodopsin and the high conservation detected in the FAR gene cluster, even across marine-freshwater transitions, point to an important role in the physiology and ecology of some of the most abundant aquatic microbes. Perhaps extending the “solar energy” strategy to deeper waters where a large part of the water column microbiome activity takes place.

## METHODS

### Screening of rhodopsin proteins from single-amplified genomes and metagenomic datasets

To evaluate the presence and the number of rhodopsin sequences in the euphotic marine microbiome, a compendium of 12,715 single-amplified genomes (SAGs) were downloaded from NCBI under BioProject PRJEB33281 (15) and subjected to study. Prior to the analysis, the degree of completeness and contamination of SAGs were estimated using CheckM v1.1.2 (61), and SAGs with > 50 % completeness and < 5 % contamination were kept. Taxonomic classification of SAGs was performed using the GTDB-Tk v 2.1.0 tool (62) using the Genome Taxonomy Database (GTDB) release R207 (63).

Metagenomes from different marine and freshwater datasets were used to retrieve similar protein sequences to FArhodopsins, as well as the contigs containing them. Marine datasets included metagenomic assemblies and PacBio CCS reads from a local time series in the Mediterranean Sea (28, 29), Hawaii Ocean Time-series (HOTs) and Bermuda Atlantic Time-series Study (BATS) (30), as well as from large oceanic expeditions, such as Tara Oceans (31) and GEOTRACES (30). Freshwater datasets represented metagenomic assemblies from Lake Baikal (37, 38), Spanish reservoirs (34), and a compendium of 17 different freshwater lakes located in Europe and Asia (36). Assembled contigs from marine and freshwater metagenomes longer or equal to 5Kb were assigned family- or genus-level classification if at least 50 % of the genes shared the same best-hit taxonomy. Contigs failing this threshold were classified at the level of phylum.

Recovery of rhodopsin proteins was achieved using a custom and curated database of rhodopsin sequences, comprising all the families described so far from both type-I and type-III rhodopsins. Hidden Markov models (HMMs) of these families were built using hmmbuild (64) after a previous alignment by muscle (65). Putative protein sequences were then screened using hmmscan (64). Only sequence hits >= 200 aa, with an evalue < 1e-10 and a bitscore >= 75 were kept.

### Phylogenetic classification of microbial rhodopsins

A maximum likelihood phylogenetic tree was built with all the type-I microbial rhodopsins recovered from the pool of marine SAGs, as well as from the metagenomic sequences retrieved from marine and freshwater datasets. As references, we included well-characterized rhodopsin proteins with several functions described so far (H^+^, Na^+^, Cl^-^ and sensory rhodopsins). Sequences were first aligned with muscle (65), and the resulting alignment trimmed with trimal (66) to remove regions with gaps in more than 80% of the sequences. The phylogenetic tree was built with iqtree (67) with 5000 ultrafast bootstraps and the -m MFP option to find the best model that fitted to our data.

### Determination of gene clusters carrying FArhodopsins

In order to determine the location of the FArhodopsin and the absence of it in other genomes from the same taxonomic group, Pelagibacterales SAGs containing the second rhodopsin were compared using blastp (68) against the isolate genomes HIMB083 (genomospecies Ia.3/V), HTCC7011 (genomospecies Ia.3/I), and HTCC1062 (genomospecies Ia.1/I). Genomic regions were then extracted and genes predicted for further analyses. In a similar approach, the genomic islands containing the rhodopsin in HIMB59 and the gammaproteobacterial *Pseudothioglobus* were determined and extracted by comparing the genomes against closely related (>90% average nucleotide identity [ANI]) SAGs.

### Taxonomic and functional annotation of rhodopsin-carrying contigs

Prodigal v2.6.3 (69) was used to predict genes from contigs retrieved from the individual SAGs, as well as from the metagenomic assemblies. Predicted protein-encoded genes were taxonomically and functionally annotated against the NCBI NR database using DIAMOND 0.9.15 (70) and against COG (71) and TIGRFAM (72) using HMMscan v3.3 (64). Domains within proteins were predicted using InterPro (73). In order to determine the location of predicted proteins within the gene cluster, transmembrane domains and signal peptides were identified with DeepTMHMM (74) and SignalP v6.0 (75), respectively.

### Retrieval of SPFH proteins and Phylogenetic tree of flotillin sequences

Domain sequences of the SPFH family of proteins were downloaded from the Pfam database (76) and then used to search for candidates within SAGs with hmmscan. The resulting flotillin sequences were then phylogenetically classified. Reference sequences from eukaryotic flotillins Flot-1 and Flot-2 (*Bos Taurus, Homo sapiens*, and *Rattus norvergicus*) together with well-characterized sequences from *Bacillus, Staphylococcus* and *E. coli* were downloaded from the Uniprot database. We also included the closest hits determined by a blastp search against the NCBI protein database. Sequences were first aligned with muscle, and the resulting alignment trimmed with trimal to remove regions with gaps in more than 80% of the sequences. A maximum-likelihood phylogenetic tree was built with iqtree with 5000 ultrafast bootstraps and the -m MFP option to find the best model that fitted to our data.

### Metagenomic and metatranscriptomic read recruitment

Illumina raw reads from Tara Oceans metagenomes and metatranscriptomes were downloaded from the European Nucleotide Archive (ENA) following the accession numbers PRJEB1787 and PRJEB6608, respectively. Eight additional samples covering a metagenomic depth profile from the Mediterranean Sea during thermal stratification (summer -- 15, 30, 45, 60, 75, 90, 1000 and 2000m) were downloaded from NCBI BioProjects PRJNA352798 and PRJNA257723. Raw reads were trimmed with Trimmomatic v0.39 (77). Coverage values were calculated by read alignment (in subsets of 20 million reads) against the genomic islands containing the FArhodopsin using BLASTN v2.9.0 (90 % identity, > 50 bp alignment). At least 70% of the length was required to be covered by reads to be considered a recruit. Reads were normalized by the size of the contig in Kb and by the size of the metagenome in Gb (RPKGs). Transcripts from metatranscriptomic samples were aligned using BLASTN (98 % identity, > 50 bp alignment) and normalized as transcripts per kilobase of gene per million reads (RPKM).

### In-silico determination of protein structures

We used AlphaFold (41), a protein structure prediction server hosted on Google Colab (78), to predict the structures of three FArhodopsins (AG-390-L03 of Pelagibacterales, AG-337-N20 of HIMB59, and AG-313-D08 of *Pseudothioglobus*) using standard parameters. We then used PyMOL (https://pymol.org/2/) to align these structures to the structure of a blue-light absorbing proteorhodopsin from Med12 (PDB ID 4JQ6). Since the FArhodopsins and proteorhodopsin shared high overall similarity and the positions of the lysine residues, which are putative binding residues for retinal coincided, we transferred the all-trans retinal molecule from proteorhodopsin (PDB ID 4JQ6) to the FArhodopsins for further analysis of the retinal binding pocket.

## ACKNOWLEDGEMENTS

This work was supported by grant “FLEX3GEN” PID2020-118052GB-I00 (cofounded with FEDER funds) from the Spanish Ministerio de Economía, Industria y Competitividad to ML-P and FR-V. JMH-M was supported with a PhD fellowship from Margarita Salas program, cofounded by the Spanish Ministerio de Universidades and the European Union -- Next Generation EU (2021/PER/00020). RS was supported by the Simons Foundation grants 510023 and 827839. The work of AA was supported by funding of the German Research Foundation to Dr. Tobias Moser via the Multiscale Bioimaging - Cluster of Excellence (EXC 2067/1-390729940).

## AUTHORS’ CONTRIBUTIONS

JMH-M conceived the study. JMH-M, ML-P, AA, EP, KK and FR-V analysed the data. JMH-M, ML-P, AA, VG, RS and FR-V contributed to write the manuscript.

## SUPPLEMENTARY MATERIAL

**Figure S1**. Complete protein sequence alignment of 6 proteorhodopsin and 6 FArhodopsin sequences. Sequences come from 6 SAGs (two of each taxa) containing both rhodopsins in their genomes. Dark and light grey regions represent conserved residues within the alignment.

**Figure S2**. Maximum-likelihood phylogenetic tree of flotillin genes retrieved from marine SAGs. Reference flotillin sequences and their accession numbers within brackets are colored in black. Sequences in red represent sequences from marine SAGs that were not found near a FArhodopsin. Sequences in blue correspond to the flotillins of the three taxa coding for the tandem flotillin-FArhodopsin. Sequences in this branch were condensed for simplicity.

**Figure S3**. Relative abundance (measured in RPKG) of the FAR gene cluster in Tara Ocean metagenomes. Each province has been divided by depth in surface (SRF), deep chlorophyll maximum (DCM) and mesopelagic (MES) regions. The bars have been colored according to the microbes (blue, *Pelagibacterales;* purple, HIMB59 and green, *Pseudothioglobus*).

**Table S1**. Summary of marine SAGs, with their taxonomic classification and the number of type-I and type-III rhodopsin genes detected.

**Table S2**. Functional annotation of genes encoded within the FAR gene clusters of Pelagibacterales, HIMB59 and *Pseudothioglobus*. Only one FAR gene cluster per taxa is shown.

**Table S3**. Number of rhodopsin sequences per metagenomic dataset.

**Table S4**. Protein alignment of rhodopsin metagenomic sequences. Amino acids involved in the ion translocation activity are colored in blue. The lysine involved in retinal binding is colored in red.

