## Supplementary material for "Flotillin-Associated rhodopsin (FArhodopsin), a widespread paralog of proteorhodopsin in aquatic bacteria with streamlined genomes": Figure S1

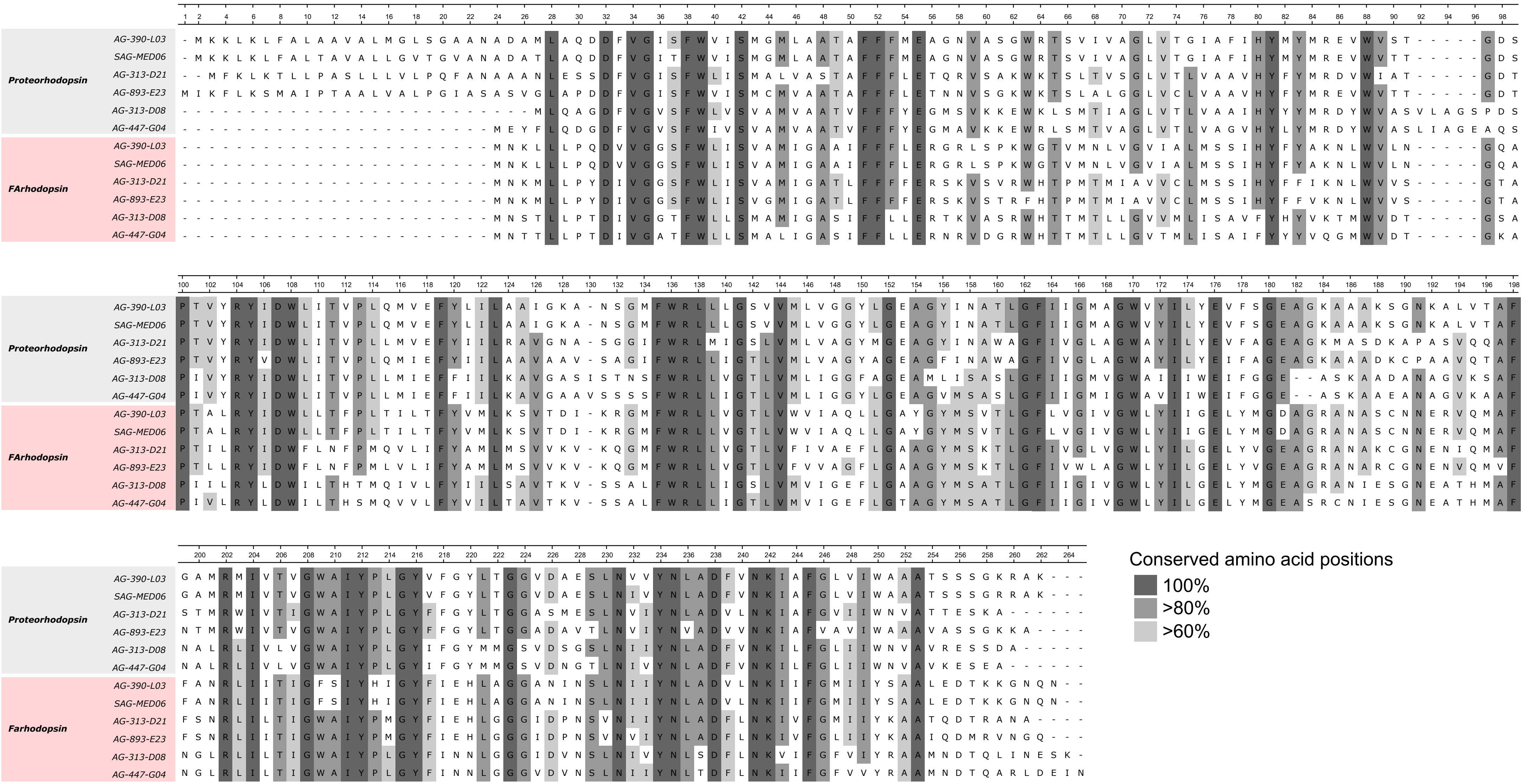


**Figure S1.** Complete protein sequence alignment of 6 proteorhodopsin and 6 FArhodopsin sequences. Sequences come from 6 SAGs (two of each taxon) containing both rhodopsins in their genomes. Dark and light grey regions represent conserved residues within the alignment.
