## Supplementary material for "Flotillin-Associated rhodopsin (FArhodopsin), a widespread paralog of proteorhodopsin in aquatic bacteria with streamlined genomes": Figure S2

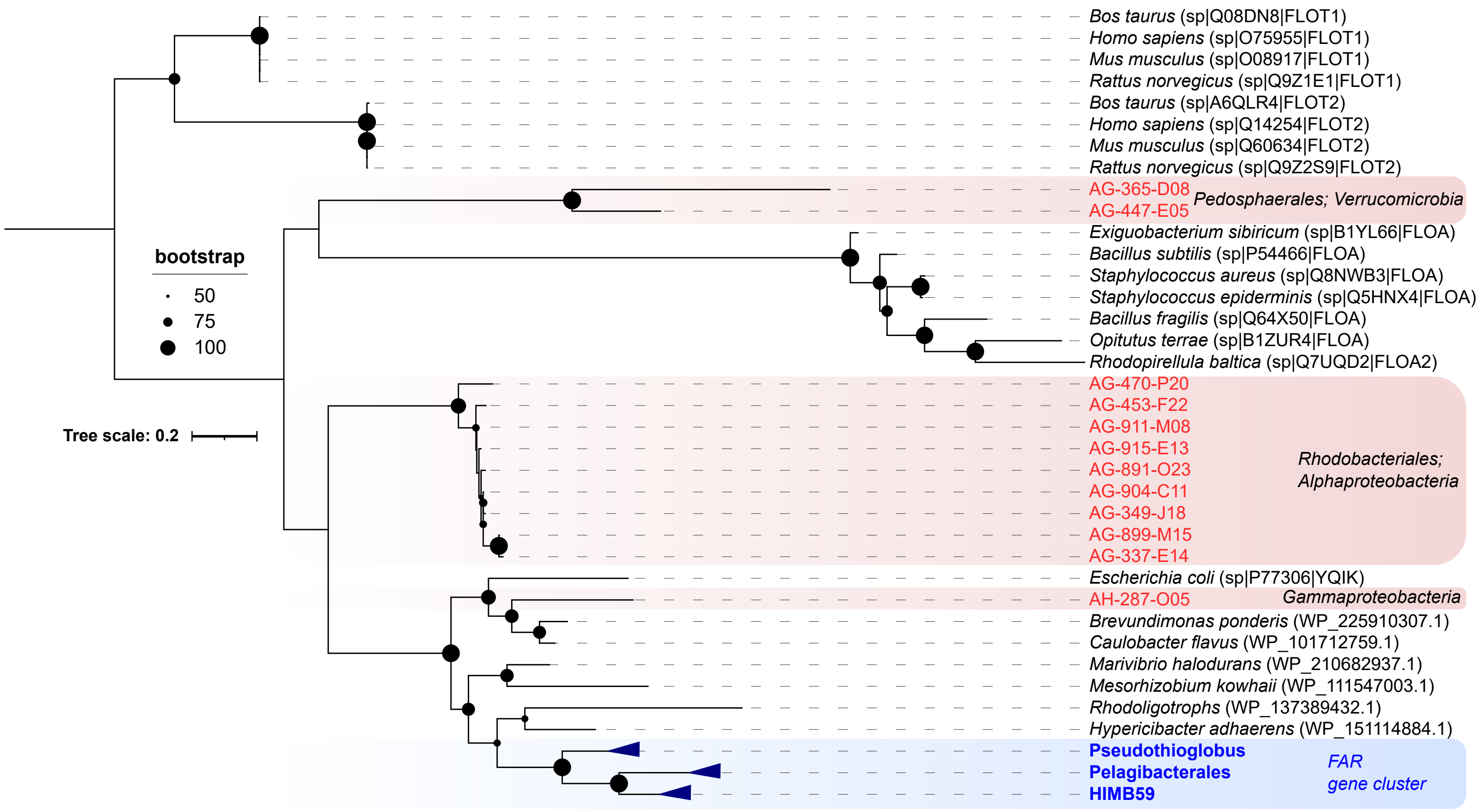


**Figure S2.** Maximum-likelihood phylogenetic tree of flotillin genes retrieved from marine SAGs. Reference flotillin sequences and their accession numbers within brackets are colored in black. Sequences in red represent sequences from marine SAGs that were not found near a FArhodopsin. Sequences in blue correspond to the flotillins of the three taxa coding for the tandem flotillin-FArhodopsin. Sequences in this branch were condensed for simplicity.
