## Supplementary material for "Flotillin-Associated rhodopsin (FArhodopsin), a widespread paralog of proteorhodopsin in aquatic bacteria with streamlined genomes": Figure S3

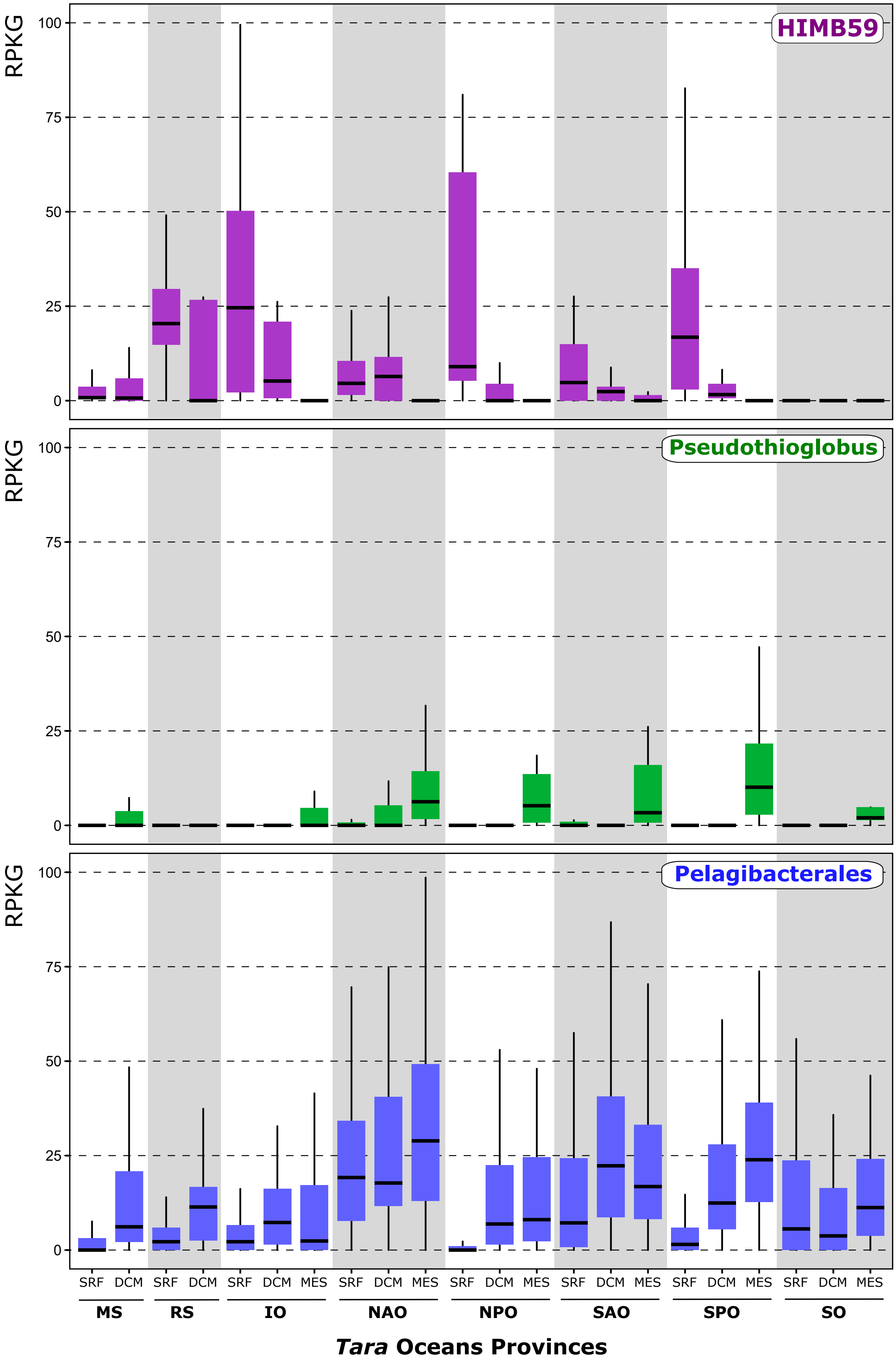


**Figure S3.** Relative abundance (measured in RPKG) of the FAR gene cluster in Tara Ocean metagenomes. Each province has been divided by depth in surface (SRF), deep chlorophyll maximum (DCM) and mesopelagic (MES) regions. The bars have been colored according to the microbes (blue, Pelagibacterales; purple, HIMB59 and green, Pseudothioglobus). MS -- Mediterranean Sea; RS -- Red Sea; IO -- Indian Ocean; NAO and SAO -- North and South Atlantic Ocean; NPO and SPO -- North and South Pacific Ocean; SO -- Southern Ocean.
